# Trypanosomal MICOS is assembled on non-respiring mitochondrial crista precursors and associates with two integral microproteins

**DOI:** 10.64898/2026.07.22.740009

**Authors:** Michala Boudová, Teresa Wagner, Tomáš Bílý, Martina Tesařová, Corinna Benz, Hassan Hashimi

## Abstract

The mitochondrial contact site and cristae organizing system (MICOS) is a multiprotein complex that shapes crista junctions and maintains inner and outer membrane contacts. MICOS coordinates the assembly of electron transport chain complexes, a prerequisite for cellular respiration. Indeed, MICOS is lost in eukaryotes that dispensed with cellular respiration, suggesting that its assembly depends on the presence of an active respiratory chain. *Trypanosoma brucei* provides a unique system to test this hypothesis as its mitochondrion undergoes developmentally regulated remodeling. In the insect stage, the mitochondrion contains cristae with an active electron transport chain, whereas the mammalian bloodstream form possesses precursor cristae with stub-like morphology that lack respiratory activity. MICOS has been characterized in the insect stage but remains unexamined in the bloodstream form. Here, we demonstrate that all MICOS subunits assemble onto precursor cristae, retaining conserved interactions with both outer and inner membrane protein machineries. This is somewhat unexpected given the co-occurrence of MICOS with active cellular respiration in nature. Furthermore, we identify novel MICOS-associated proteins that are dispensable for its stability, suggesting auxiliary rather than core roles in MICOS function. Together, our findings establish that MICOS assembly precedes cellular respiratory competence and expand its interaction landscape in trypanosomatids.

## Introduction

Mitochondria are organelles of endosymbiotic origin, best known for their role in energy production, and therefore often referred to as the ‘powerhouse of the cell’. As a vestige of their ancestry, mitochondria are surrounded by a double membrane, that delineates distinct sub-compartments, enabling spatial separation of functions. The outer mitochondrial membrane (OMM) encapsulates the organelle, while the mitochondrial inner membrane (IMM) acts as a highly impermeable barrier that sequesters the mitochondrial matrix from the intermembrane space (IMS) (Mukherjee et al., 2021).

The IMM is typically folded into numerous invaginations known as cristae. The crista membrane (CM) is enriched in respiratory chain complexes that pump protons from the matrix into the crista lumen, thereby generating the electrochemical gradient that fuels ATP synthesis through oxidative phosphorylation (OXPHOS)(Colina-Tenorio et al., 2020; Pánek et al., 2020). These structures are connected to the inner boundary membrane by constrictions at their base, termed crista junctions (CJs), which restrict diffusion between the crista lumen and the IMS. The assembly and maintenance of CJs is facilitated by the Mitochondrial Contact Site and Cristae Organizing System (MICOS), which also participates in the formation of contact sites between the outer and the inner mitochondrial membranes (Fig. 1A; Daumke & van der Laan, 2025; Mukherjee et al., 2021; van der Laan et al., 2016).

**Fig. 1.**
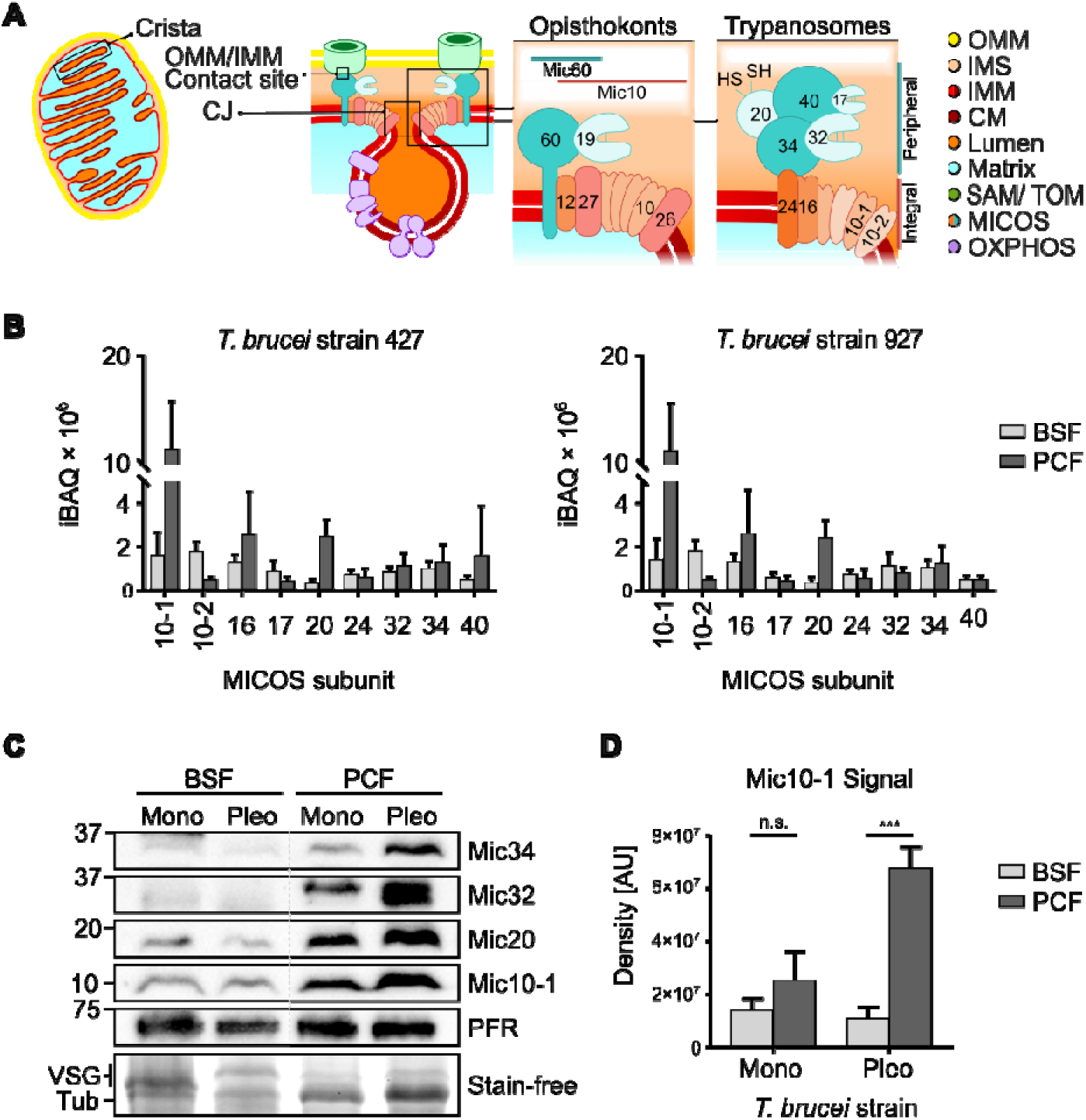
MICOS subunits are expressed in *T. brucei* BSF with precursor cristae. **(A)** Schematic overview of mitochondrial architecture and MICOS organization. The leftmost diagram illustrates the overall mitochondrial structure (adapted from Pánek et al., 2020), with subcompartments color-coded as indicated in the legend on the right. The inner mitochondrial membrane (IMM) is subdivided into the inner boundary membrane (IBM) and the crista membrane (CM), highlighted by darker shading within the crista lumen. The cristae along with the CJs and the OMM/IMM contact sites are highlighted in boxes and enlarged in the following scheme. The two rightmost diagrams (adapted from Eichenberger et al., 2019) compare MICOS organization in Opisthokonts (e.g. yeast) and trypanosomes. In yeast, the integral and peripheral MICOS subcomplexes are arranged laterally, whereas in trypanosomes the membrane-peripheral module is positioned vertically relative to the integral one. Individual MICOS subunits are indicated by the terminal digits of their names. **(B)** Abundance of MICOS subunits across bloodstream (BSF) and procyclic (PCF) life cycle stages of *T. brucei* laboratory strains 927 and 427, as determined by LC-MS/MS (Tinti & Ferguson, 2022). Bars represent mean iBAQ values; error bars indicate standard deviation. **(C)** Immunoblot analysis of MICOS subunit levels in BSF and PCF stages of monomorphic (Mono) or pleomorphic (Pleo) *T. brucei* strains. Molecular weight is shown on the left, antibodies are indicated on the right. PFR serves as loading control. Stain-free gel highlights the disappearance of the major BSF surface protein VSG in PCF samples. **(D)** Quantification of Mic10-1 protein levels based on the band density on immunoblots; bars represent mean values and error bars indicate standard deviation (n=4). Statistical significance was determined using an unpaired two-tailed Welch’s t-test (p < 0.0001).

In Opisthokonts, the eukaryotic clade that encompasses animals and yeast, MICOS consists of at least six subunits organized into two adjacent integral membrane subcomplexes in a horizontal arrangement (Fig. 1A). Each subcomplex is assembled around a respective core subunit, Mic10 or Mic60. The Mic10-containing module reinforces membrane bending at CJs (Barbot et al., 2015; Bohnert et al., 2015) and contributes to the biogenesis of lamellar cristae in vertebrates (Stephan et al., 2020). The Mic60-containing subcomplex is proposed to constrict CJs (Bock-Bierbaum et al., 2022) and mediates contacts with protein complexes of the OMM, including the β-barrel insertase Sam50 (Huynen et al., 2016; Tang et al., 2020). Through this interaction, Mic60 also supports protein import mediated by the mitochondrial intermembrane space assembly (MIA) pathway (Eichenberger et al., 2019; Rampelt et al., 2022; van der Laan et al., 2016). Importantly, the stability of individual MICOS subunits is highly interdependent, such that loss of one component often destabilizes several others (Harner et al., 2011; Hoppins et al., 2011). Additionally, MICOS is also involved in the biogenesis of respiratory chain complexes III and IV within the CM (Colina-Tenorio et al., 2026; Zerbes et al., 2025).

Positioned at the core of MICOS, the key subunit Mic60 is conserved from extant Alphaproteobacteria to Opisthokonts, with the highest degree of conservation in its C-terminal mitofilin domain, which is essential for membrane remodelling (Bock-Bierbaum et al., 2022; Huynen et al., 2016; Muñoz-Gómez et al., 2023). However, the MICOS complex is highly divergent in the phylum Euglenozoa, where it consists of nine subunits organized into an integral membrane module and a peripherally-associated subcomplex located within the IMS. In contrast to the lateral arrangement of MICOS modules in Opisthokonts, these two subcomplexes are most likely oriented perpendicular to the plane of the IMM (Fig. 1A; Eichenberger et al., 2019). In the phylum Kinetoplastea that contains trypanosomatids, the integral module is organized around the two Mic10 paralogs. Moreover, the canonical Mic60 has been replaced by two cryptic members of the mitofilin family, Mic34 and Mic40, which likely form the core of the peripheral subcomplex (Eichenberger et al., 2019; Sheikh et al., 2025). Another component of this module is the thioredoxin-like protein Mic20, which appears to be directly involved in a yet undefined aspect of the MIA pathway (Kaurov et al., 2022; Sheikh et al., 2025).

Apart from the MICOS complex, which promotes negative curvature at CJs, ATP synthase dimers shape cristae by oligomerizing at crista rims, inducing positive membrane curvature (Mukherjee et al., 2021; van der Laan et al., 2016). Despite these apparently antagonistic effects on crista architecture, a fraction of Mic10 has been shown to interact with ATP synthase, coordinating the activities of these two complexes (Cadena et al., 2021; Rampelt et al., 2022).

To meet changing energy demands, cristae undergo continuous remodelling in response to physiological or developmental cues (Mukherjee et al., 2021; Pánek et al., 2020). This phenomenon is also observed in *Trypanosoma brucei*, a digenetic unicellular parasite and a well-established model organism. As it alternates between mammalian and insect hosts, its single mitochondrion undergoes striking ultrastructural and metabolic reorganization (Bílý et al., 2021; Zíková, 2022). In bloodstream forms (BSF), parasites reside in a glucose-rich environment, and thus energy production occurs by substrate-level phosphorylation. Consequently, the mitochondrion is highly reduced, adopting a tubular morphology with sparse, stub-like cristae that do not contain any electron transport chain (ETC) complexes. We term these structures precursor cristae. Following transmission to the insect vector, parasites differentiate into the procyclic form (PCF) adapted to the glucose-poor insect midgut, leading to expansion of the mitochondrion into a reticulated network replete with mature discoidal cristae and concomitant activation of OXPHOS (Bílý et al., 2021; Zíková, 2022). While the MICOS complex has been characterized in PCF (Kaurov et al., 2018; Sheikh et al., 2025), its presence, organization and function in BSF are addressed here for the first time. Our findings demonstrate that MICOS assembly occurs in the absence of an active respiratory chain, which is unexpected, given that anaerobic eukaryotes have lost MICOS concomitantly with the loss of mitochondrial respiration (Lewis et al., 2020; Pánek et al., 2020). Furthermore, our analyses have uncovered novel MICOS interactors, revealing previously unrecognized aspects of MICOS organization.

## Results

### MICOS subunits are expressed in cells with precursor cristae

Given the low occupancy of cristae in BSF as compared to PCF (Bílý et al., 2021), we hypothesized a corresponding downregulation of MICOS expression. To assess MICOS abundance across life cycle stages, we reanalyzed a previously published liquid chromatography-tandem mass spectroscopy (LC-MS/MS) dataset (Tinti & Ferguson, 2022). The hypothesis is valid for Mic10-1 and Mic20 (Fig. 1B). Surprisingly, in BSF of strain 427, Mic10-2 and Mic17 were upregulated, whereas all other subunits showed modest upregulation in PCF.

We next validated these proteomic results by using antibodies recognizing the subunits Mic10-1, Mic20, Mic32, and Mic34 to assay their levels in PCF and BSF lysates derived from equal numbers of whole cells (Fig. 1C). In addition to the monomorphic PCF and BSF cell lines with TREU927/4 and Lister 427 genetic backgrounds, respectively, which are unable to complete their life cycle, we also analyzed lysates from both life cycle stages of a pleomorphic cell line that retains the capacity for differentiation and the associated mitochondrial remodeling (Cai et al., 2021; Overath et al., 1986). This allowed us to follow MICOS regulation during crista maturation as parasites transition from BSF to PCF. Immunoblot analyses detected all examined MICOS subunits in both life cycle stages, although Mic32 and Mic34 were only weakly detected in BSF samples compared to Mic10-1 and Mic20. Overall, the observed trends were partially consistent with previously published data (Tinti & Ferguson, 2022), which showed more pronounced Mic10-1 and Mic20 upregulation compared to Mic32 and Mic34 (Fig. 1C). Notably, in the pleomorphic line, upregulation of all examined subunits in PCF appeared more pronounced relative to their monomorphic counterparts (Fig 1D). In conclusion, we established that MICOS subunits are present in mitochondria with precursor cristae and that most subunits are upregulated during cristae maturation.

### The mitofilin family Mic34 co-localizes with precursor cristae

To facilitate biochemical characterization of MICOS in BSF, we generated monomorphic cell lines in which Mic10-1, Mic10-2, Mic34, and Mic40 were endogenously C-terminally tagged with an HA epitope (Fig. S1A). We next assessed mitochondrial targeting of the tagged proteins. All subunits separated into the mitochondria-enriched fraction upon digitonin fractionation of *T. brucei* cells (Fig. 2A). Mitochondrial localization was further verified by indirect immunofluorescence assay, with no signal observed in the parental cell line, confirming the specificity of HA detection (Fig. 2B, S1B). Sub-organellar distribution of Mic34-HA was resolved by immunoelectron microscopy. This task was complicated by the low contrast of crista membrane staining resulting from sample processing for immunogold labelling (Tokuyasu, 1978), the diminutive size of BSF cristae (Bílý et al., 2021) and the fact that only HA-epitopes exposed on the surface of the thick-section are accessible to antibody. Nevertheless, we identified four independent mitochondrial immunogold signals. In all cases, these signals were located adjacent to the internal looped membranes, allowing us to conclude with reasonable confidence that Mic34-HA associates with precursor cristae (Fig. 2C, S2; Movie S1-4). Based on this finding, we hypothesized that the MICOS complex is assembled on precursor cristae.

**Fig. 2.**
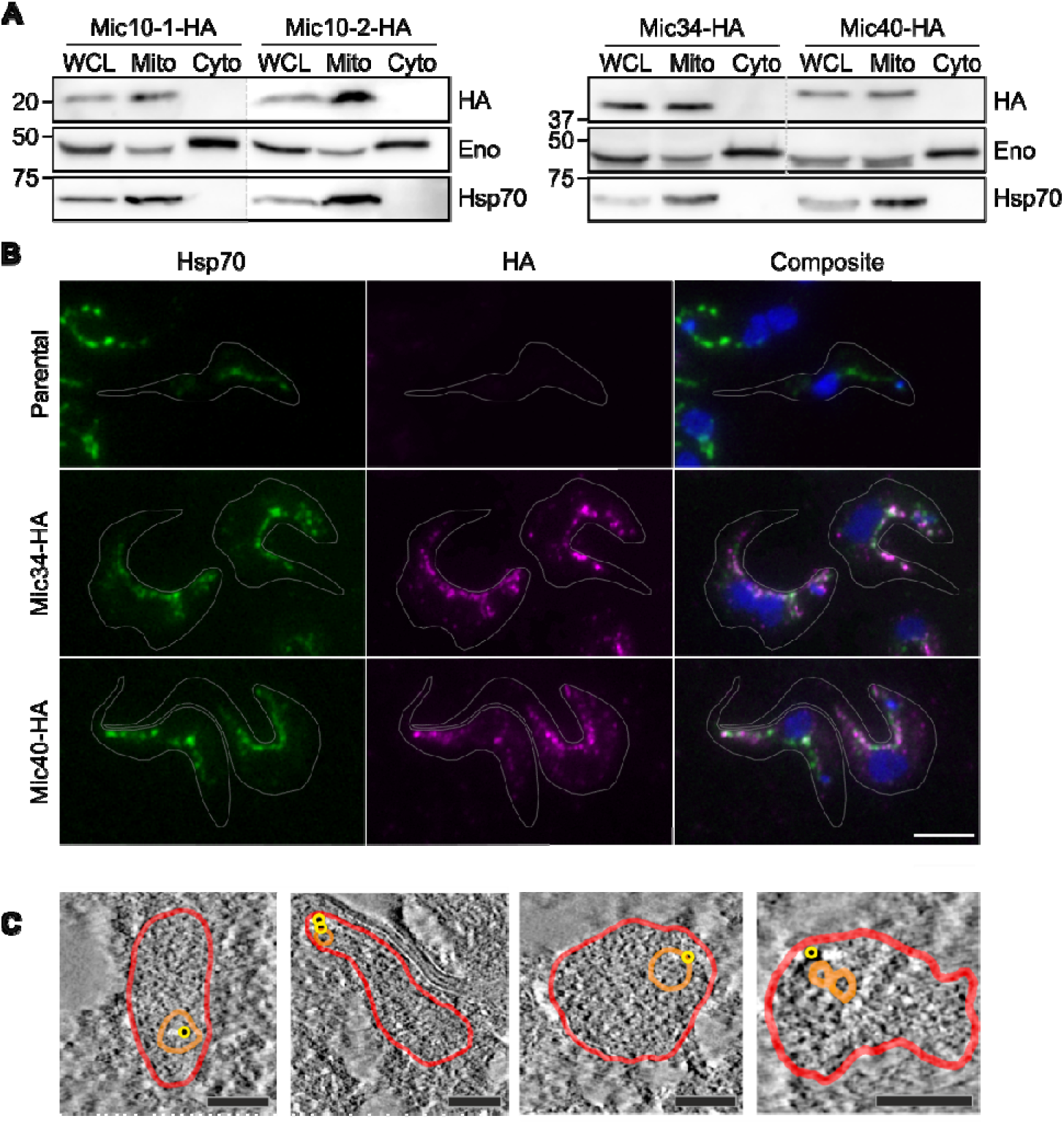
MICOS subunits localize to mitochondria containing precursor cristae. **(A)** Immunoblot analysis of MICOS subunit localization following selective permeabilization with digitonin to separate organellar (Mito) and cytosolic (Cyto) subcellular fractions. Fractions were analyzed alongside whole-cell lysate (WCL). The respective cell lines are indicated above the blot, molecular weight on the left, and antibodies on the right side. Enolase (Eno) served as the cytosolic marker and heat shock protein (Hsp70) as the mitochondrial marker. **(B)** Immunofluorescence assay for selected cell lines corroborates the results shown in (A). Cell lines are indicated on the left; the parental line was processed in parallel as a negative control. Antibodies are indicated above the panel; Hsp70 marks mitochondria, and DAPI (blue) stains DNA. The composite image demonstrates substantial overlap of the fluorescence signals. Scale bar, 5 μm. **(C)** Electron tomograms of immunogold-labeled Mic34-HA. Mitochondrial membranes are traced in red, the crista membrane in orange, and gold nanoparticles are highlighted with yellow circles. Scale bar, 100 nm.

### The complete MICOS complex assembles onto precursor cristae and engages in extra-MICOS interactions

To determine whether MICOS subunits assemble into a higher-order complex in BSF, we first analyzed detergent-solubilized mitochondrial fractions by blue native PAGE (BN-PAGE). All HA-tagged MICOS subunits migrated within a high-molecular-weight assembly corresponding to the endogenous MICOS complex detected using anti-Mic10-1 antibody (Fig. 3A, S1). These data demonstrate that MICOS subunits are incorporated into a common macromolecular complex despite the absence of a fully developed respiratory chain and mature discoidal cristae in BSF.

**Fig. 3.**
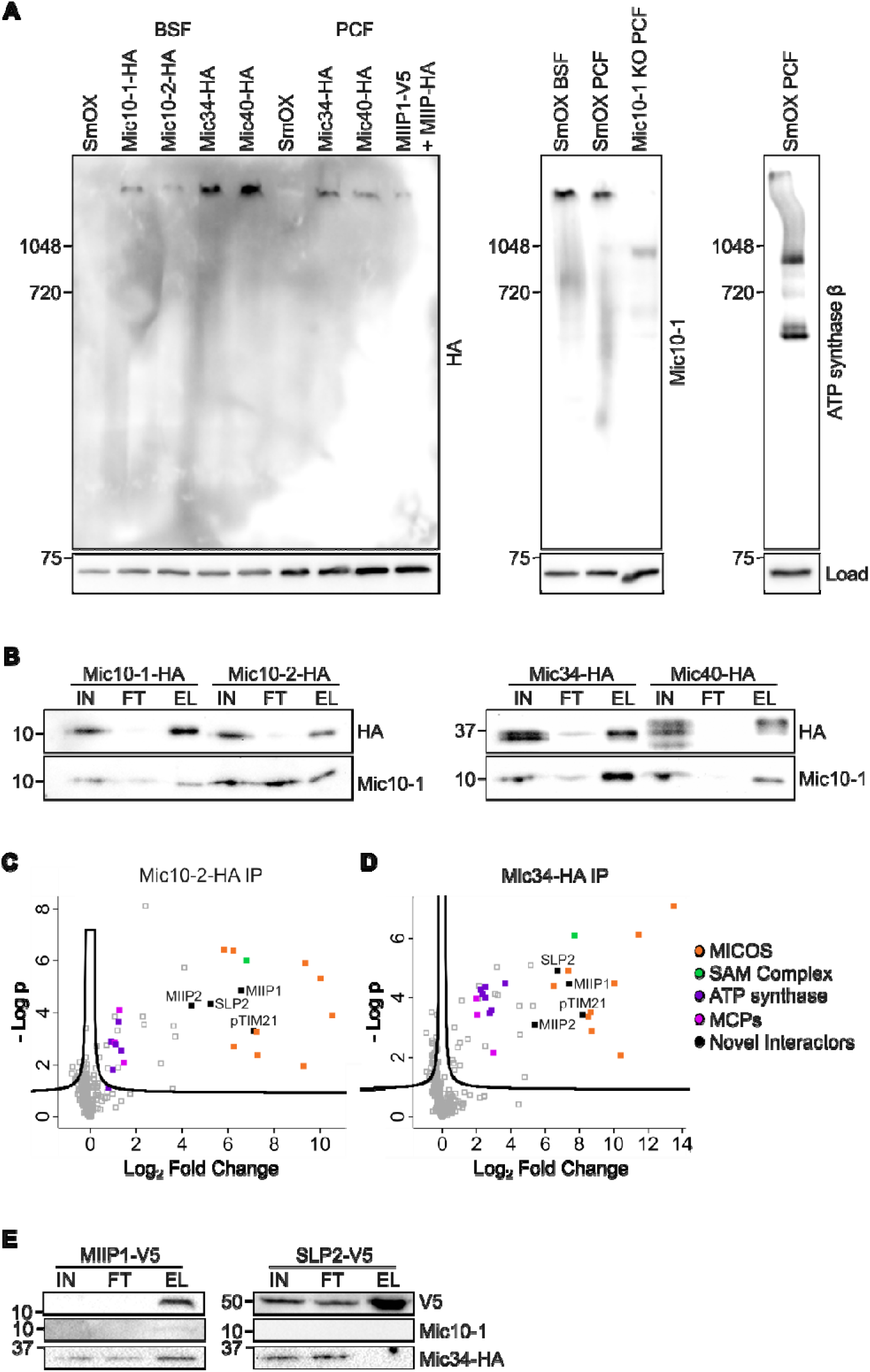
The complete MICOS complex assembles on precursor cristae. **(A) Left:** Blue native PAGE analysis of BSF cell lines expressing HA-tagged Mic10-1, Mic10-2, Mic34, or Mic40 followed by immunoblotting with anti-HA antibody. PCF cell lines expressing HA-tagged Mic34 or Mic40 were included as positive controls. In addition, PCF dual-tagged MIIP1-V5+MIIP2-HA was included. All tagged MICOS subunits migrate within a high-molecular weight complex. **Middle:** Endogenous MICOS complex in both BSF and PCF detected with anti-Mic10-1 antibody. PCF Mic10-1 knock-out (KO) cell line served as negative control. **Right:** Detection of ATP synthase subunit β as an independent high-molecular-weight mitochondrial complex. An aliquot of each sample was denatured and analyzed by SDS-PAGE shown below. Immunodetection of Hsp70 served as a loading control (Load). Molecular weight is indicated on the left and antibodies on the right. (**B**) Immunoblot analysis to verify immunoprecipitation (IP) using the indicated HA-tagged MICOS subunits as baits and the co-enrichment of Mic10-1. Protein enrichment was monitored throughout the procedure, from input (IN) to flow-through (FT) and eluate (EL), the latter representing the bead-bound material. Molecular weight is indicated on the left and antibodies on the right. **(C, D)** Volcano plots depicting protein enrichment in (C) Mic10-2-HA and (D) Mic34-HA IPs. The x-axis shows log_2_-transformed fold change relative to the parental cell line, and the y-axis represents −log-transformed P-values from triplicate measurements. Enriched proteins are grouped into color-coded categories, as indicated in the legend on the right. Four novel interactors identified in both IPs are labelled. **(E)** Immunoblot analysis of reciprocal IPs for two candidate interactors identified in (C-D): MIIP1-V5 and SLP2-V5. Cell lines and fractions (labelled as in (B)) are indicated above the blot, molecular weight on the left, and antibodies on the right.

We therefore proceeded to characterize the composition of this assembly and identify potential interaction partners. To this end, all four HA-tagged cell lines were subjected to immunoprecipitation (IP) *via* their epitope tags. Mic10-1 co-enrichment was detected in all samples (Fig. 3B), suggesting that the entire MICOS complex was recovered. To validate this hypothesis, Mic10-2-HA and Mic34-HA immunoprecipitates were examined by LC-MS/MS as representatives of the Mic10 and the mitofilin subcomplexes respectively (Eichenberger et al., 2019). As shown in Fig. 3C-D, all nine MICOS subunits were detected in both IPs, confirming that the complex is present on precursor cristae.

Furthermore, extra-MICOS interactions previously observed in PCF (Cadena et al., 2021; Kaurov et al., 2018) were also detected in BSF. The OMM β-barrel insertase Sam50 was enriched in both experiments. Mic34-HA additionally immunocaptured F_O_F_1_-ATP synthase subunits from both F_O_ and F_1_ moieties (Fig. 3D; Table S1). The comparatively lower enrichment of F_O_F_1_-ATP synthase subunits in Mic10-2-HA IP (Fig. 3C) is consistent with such a result in PCF (Cadena et al., 2021).

Novel extra-MICOS interactions were also identified. Similar to the F_O_F_1_-ATP synthase subunits, mitochondrial carrier proteins (MCPs) were preferentially enriched in the Mic34-HA IP (Fig. 3D; Table S1). Among these are MCP5, the ADP/ATP carrier(Gnipová et al., 2015; Peña-Diaz et al., 2012), MCP12, a dicarboxylate/ tricarboxylate antiporter (Colasante et al., 2018), and MCP13, a putative GDP/GTP carrier (Colasante et al., 2009).

Finally, four additional proteins were reproducibly detected in both IPs (Fig. 3C-D, Table S2), potentially constituting stable MICOS interactors. These include stromatin-like protein 2 (SLP2), previously studied in *T. brucei* (Serricchio & Bütikofer, 2021), and a putative homolog of mitochondrial inner membrane translocase subunit pTim21, identified through structural homology searches (Fig. S3). The remaining two qualify as microproteins as they are <100 amino acids long (Hassel et al., 2023) and contain a single transmembrane domain (TMD). Accordingly, we termed them MICOS-interacting integral protein (MIIP) 1 and 2.

Reciprocal IP experiments demonstrated weak enrichment of Mic10-1 and modest enrichment of Mic34-HA with MIIP1-V5, whereas no such interaction was detected with the SLP2-V5 as bait (Fig. 3E). Unfortunately, neither MIIP2 nor pTim21 could be successfully tagged in BSF, which prompted us to investigate whether these four proteins interact with Mic10-1 in PCF. Although SLP2 and pTim21 were successfully V5-tagged in this life cycle stage, neither co-immunoprecipitated Mic10-1 (Fig. S4).

Collectively, these data demonstrate that MICOS is assembled in the presence of precursor cristae and forms both conserved and novel protein interactions in BSF. Thus, we continued with our investigation of MIIP1 and MIIP2 in PCF.

### MICOS associates with two novel integral-membrane microproteins

Both microproteins were C-terminally V5-tagged in PCF (Fig. S5A), and immunofluorescence assay confirmed their mitochondrial targeting (Fig. 4A), consistent with previous reports (Moloney et al., 2023; Pyrih et al., 2023) (Table S3). Co-IP experiments demonstrated that both MIIP1-V5 and MIIP2-V5 interact with Mic10-1 (Fig. 4B), leading to the conclusion that both MIIPs also associate with MICOS in PCF. Moreover, HA-tagged MIIP2 co-immunoprecipitated MIIP1-V5, showing strong interaction between the two microproteins (Fig. 4C).

**Fig. 4.**
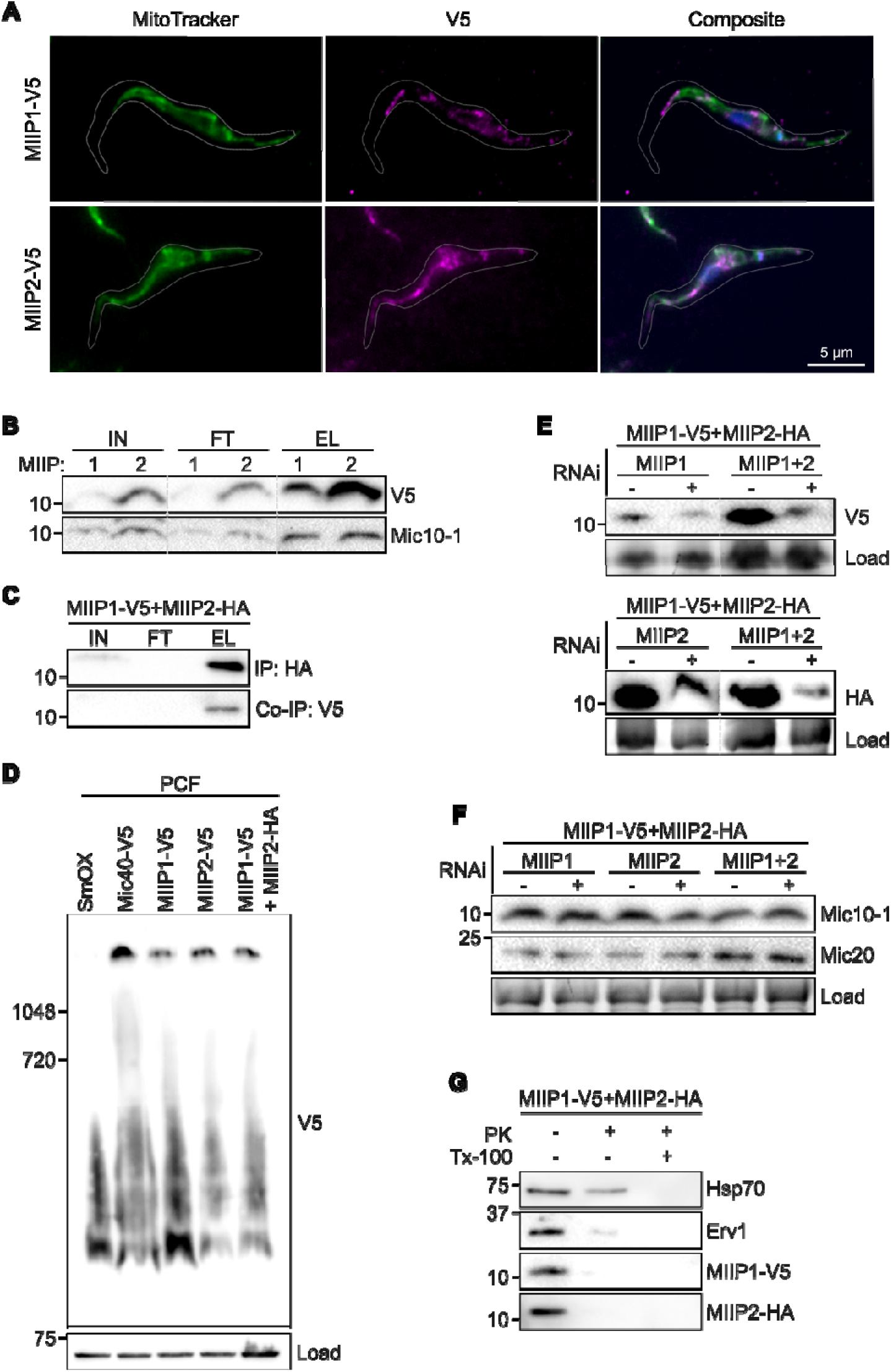
MICOS associates with two novel microproteins. **(A)** Immunofluorescence assay of MIIP1-V5 and MIIP2-V5 in PCF. Cell lines are indicated above the panel. The detection method is indicated on the left. The composite image shows overlap of the V5 and mitochondrial signals, consistent with mitochondrial localization. Scale bar, 5 μm. **(B)** Immunoblot analysis of immunoprecipitation (IP) experiments using the indicated V5-tagged candidate interactors as baits and assessing Mic10-1 co-IP. Protein enrichment was monitored across input (IN), flow-through (FT), and eluate (EL) fractions. **(C)** Immunoblot analysis of MIIP2-HA IP. Protein enrichment and labelling as in (B). (**D**) Blue native PAGE analysis of PCF cell lines expressing MIIP1-V5, MIIP2-V5, or dual-tagged MIIP1-V5+MIIP2-HA. PCF Mic40-V5 cell line was included as a positive control. Immunoblotting with anti-V5 antibody shows that both MIIPs migrate within a high-molecular-weight assembly consistent with MICOS. An aliquot of each sample was denatured and analyzed by SDS-PAGE shown below. Immunodetection of Hsp70 served as a loading control (Load). Molecular weight markers are indicated on the left and the antibodies on the right. **(E)** Immunoblots verifying inducible RNAi-mediated downregulation of MIIP1 and MIIP2, as indicated above the blot. **(F)** Immunoblot monitoring the effect of MIIPs depletion on the steady-state levels of representative MICOS subunits. Cell lines are indicated above. **(G)** Proteinase K protection assay performed on mitoplasts to assess the topology of MIIPs. As indicated above the immunoblot, samples were either left untreated, incubated with Proteinase K to degrade IMS-exposed proteins, or additionally treated with Triton X-100 (Tx-100) to allow degradation of matrix proteins. Erv1 and Hsp70 served as IMS and matrix controls, respectively.

To determine whether these interactions reflect stable association with MICOS, we analyzed detergent-solubilized mitochondrial fractions by BN-PAGE. MIIP1-V5, MIIP2-V5 as well as MIIP2-HA all migrated within a high-molecular-weight assembly corresponding to the endogenous MICOS complex (Fig. 3A, 4D, S5A), indicating that both microproteins are stably associated with MICOS.

To investigate the role of MIIP1 and MIIP2 in MICOS organization, we generated inducible RNAi cell lines targeting each *miip* transcript. In a dual-tagged cell line expressing MIIP1-V5 and MIIP2-HA, both proteins were simultaneously downregulated. RNAi-mediated silencing of the MIIPs, either individually or in combination, did not influence Mic10-1 or Mic20 steady state levels (Fig. 4E-F), nor *T. brucei* growth (Fig. S5B). These results are in agreement with the essential role of Mic20 in parasite growth (Kaurov et al., 2018, 2022). Notably, disruption of either MICOS subcomplex typically results in degradation of its constituent proteins (Eichenberger et al., 2019), hence the persistence of both MICOS subcomplex markers upon MIIPs downregulation is inconsistent with the behavior expected for *bona fide* MICOS subunits.

To gain further insight into their properties, we performed *in silico* analysis of MIIP1 and MIIP2. Based on our results, the two proteins are not homologs, MIIP1 is encoded by a kinetoplastid core gene (Kissinger, 2006) present in all sampled kinetoplastid genomes, whereas MIIP2 is exclusive to *T. brucei* (Table S3).

Despite their independent origins, both proteins share several noteworthy features in addition to their size. Both are predicted to fold into an extended α-helix split into two parts (Fig. 5A-B). A predicted TMD starting 4-5 amino acids from the N-terminus anchors each protein in the IMM, while the remaining two-thirds of the helix projects into the IMS and displays distinct charged and hydrophobic patches (Fig. 5A, 5C). Both trypanosomatid MIIP1 orthologues possess a four-amino-acid N-terminal motif that is predicted to function as a cleavable mitochondrial targeting sequence (MTS) by two algorithms (Table S3). However, this prediction should be interpreted with caution, as this would represent an unusually short MTS with a net charge of only +1. In contrast, MIIP2 contains a five-amino-acid N-terminal motif preceding the TMD that includes two basic residues (Fig. 5A).

**Fig. 5.**
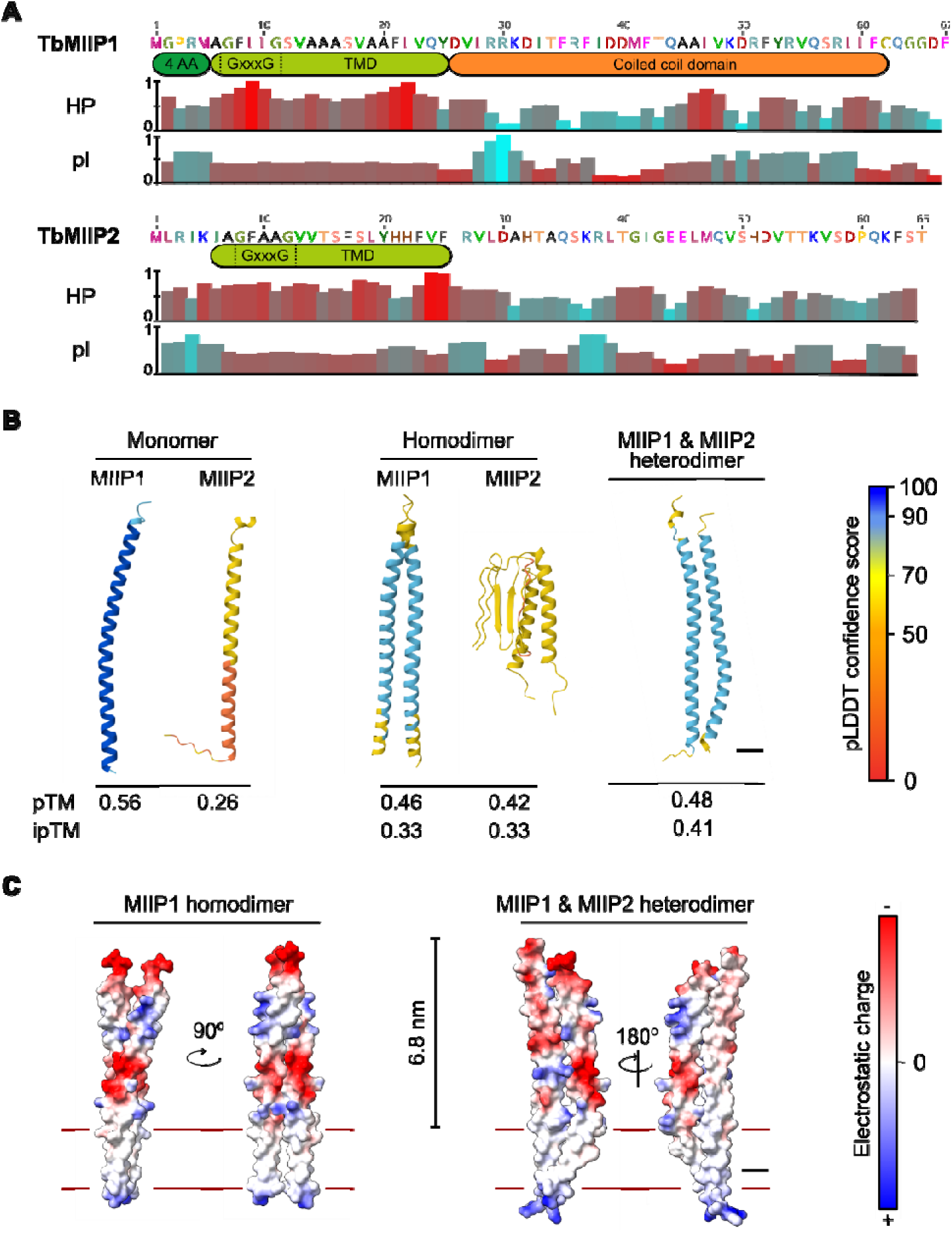
MIIP1 and MIIP2 share similar predicted biophysical properties, domain architecture and structures. **(A)** Schematic representation of predicted domains in MIIP1 and MIIP2, shown in boxes beneath sequence. The conserved GxxxG motif within each transmembrane domain (TMD) is indicated. The conserved four amino acids (4 AA) represent predicted MTS. Further below, histograms display the mean hydrophobicity (HP) and isoelectric point (pI) calculated using a 3-amino-acid sliding window. Scores range from 0 to 1, with 1 corresponding to maximal hydrophobicity (also indicated by increasing reddish hue of bin) and positive charge (increasing turquoise hue), respectively. **(B)** AlphaFold3-predicted structures of MIIP1 and MIIP2 monomers, homodimers and heterodimer. Ribbon representations are colored by confidence scores (pLDDT) according to the key on the right. Numbers below each model are their respective AlphaFold3 pTM and ipTM (for dimers) scores. Scale bar, 10 Å. **(C)** AlphaFold3-predicted MIIP1 homodimer and MIIP1/2 heterodimer are depicted as space-filling models with surface electrostatic charge mapped according to the key on the right. Double lines delineate the approximate position of the inner mitochondrial membrane. Scale bar, 10 Å.

The structural predictions suggest that the bulk of both proteins extends into the IMS. To experimentally validate this topology, we performed a proteinase K shaving assay on mitoplasts (*i.e.* mitochondria with permeabilized OMM, exposing IMS to proteolysis; Fig. 4G). As expected, the IMS protein Erv1 was degraded, whereas the matrix protein Hsp70 remained largely protected until detergent was added, supporting the validity of the assay. The C-terminal epitope tags of both MIIP1 and MIIP2 were strongly depleted by proteinase K treatment, consistent with exposure of this part of the proteins to the IMS.

The IMS extension of MIIP1 likely forms a coiled coil (CC) domain based on two lines of evidence. First, over half of the kinetoplastid orthologs are predicted to contain a CC in this region (Fig. S6, Table S4). Second, iterative hidden Markov model search using the multi-sequence alignment of MIIP1 (Fig. S6) revealed remote homology to the CC of t-SNARE, a protein involved in vesicle fusion (Dun et al., 2010). In contrast, the IMS extension of MIIP2 lacks recognizable structural motifs.

Given that CC domains as well as GxxxG motifs, present within the TMD of both MIIPs, often mediate protein interactions (Kleiger et al., 2002; Moutevelis & Woolfson, 2009), we also explored potential oligomerization. AlphaFold3 predicts MIIP1 and MIIP2 homodimerization with equally low confidence (see pTM and ipTM scores in Fig. 5B) and, moreover, the theoretical MIIP2 homodimerization substantially alters the monomeric structure. In contrast, heterodimerization between MIIP1 and MIIP2 yields a higher confidence model, with MIIP2 reaching its highest confidence score (pLDDT) in this configuration. This is consistent with the detection of MIIP1-V5 in MIIP2-HA immunoprecipitates (Fig. 4C). A space-filling rendering indicates that both the MIIP1 homodimer and MIIP1/2 heterodimer may extend about 7 nm from the membrane and display patches of negative and positive charge (Fig. 5C). Thus, a MIIP1/2 heterooligomer (Fig. 4C) and/or homooligomer of each may stably interact with components of the MICOS complex.

In conclusion, *T. brucei* MICOS associates with two integral microproteins that are dispensable for MICOS stability, arguing against their classification as *bona fide* MICOS subunits. Both proteins share similar topology, with C-terminal extensions into the IMS. However, only MIIP1 is conserved across Kinetoplastea, whereas MIIP2 is restricted to *T. brucei*.

## Discussion

Since its discovery, MICOS has been recognized as the key determinant of crista junction maintenance, forming the diffusion barrier that separates the crista membrane and lumen from the inner boundary membrane and the IMS (Colina-Tenorio et al., 2020). Recently, MICOS has been implicated in the biogenesis of respiratory chain Complex III (Zerbes et al., 2025) and Complex IV (Colina-Tenorio et al., 2026), extending its relevance beyond membrane architecture to the assembly of the machinery required for proton motive force generation. Consistently, MICOS is absent from eukaryotes that have lost the ETC during adaptation to an anaerobic lifestyle (Lewis et al., 2020; Pánek et al., 2020). Thus, our finding that MICOS assembles on precursor cristae lacking an active ETC is particularly striking.

Our IP results suggest that several molecular interactions are more robust in BSF, presumably reflecting the confined structure of precursor cristae compared to mature discoidal cristae of PCF (Bílý et al., 2021). The stronger enrichment of F_O_F_1_-ATP synthase subunits and MCPs in the Mic34 IP could indicate closer proximity and/or more frequent association to MICOS within these restricted membrane domains. One of the identified MCPs is the ATP/ADP carrier, although it is tempting to speculate that this carrier associates with F_O_F_1_-ATP synthase, there is currently no evidence for direct physical (Gnipová et al., 2015) or functional coupling (Taleva et al., 2023). These MCPs may thus reside near MICOS and/or precursor cristae in BSF for still enigmatic reasons. Furthermore, it remains unclear whether F_O_F_1_-ATP synthase dimers reside on precursor cristae (Zíková, 2022).

In contrast to equivalent IPs in PCF (Cadena et al., 2021; Kaurov et al., 2018), both MICOS IPs in this study strongly enriched Sam50. In mammals, Sam50/Mic60 interactions maintain contact sites between the outer and the IMM (Ding et al., 2015; Ott et al., 2012; Tang et al., 2020), this mechanism occurs even in extant Alphaproteobacteria *via* the Sam50 ortholog, BamA (Muñoz-Gómez et al., 2023). Given the established presence of Mic60 at crista junctions (Bock-Bierbaum et al., 2022; Stephan et al., 2020), these contact sites have been proposed to serve as fulcrums for crista membrane expansion (Tang et al., 2020).

Thus, we propose that the enhanced MICOS association with factors such as Sam50 and F_O_F_1_-ATP synthase primes precursor cristae for subsequent maturation during *T. brucei* differentiation from BSF to PCF (Bílý et al., 2021). In this hypothetical model, robust Sam50/MICOS contacts stabilize regions of membrane bending as stub-like cristae expand into discoidal shapes (Fig. 6). Concurrently, F_O_F_1_-ATP synthase dimers serve as nucleation sites for positive curvature, facilitating the transition toward mature crista architecture (Gahura et al., 2022).

**Fig. 6.**
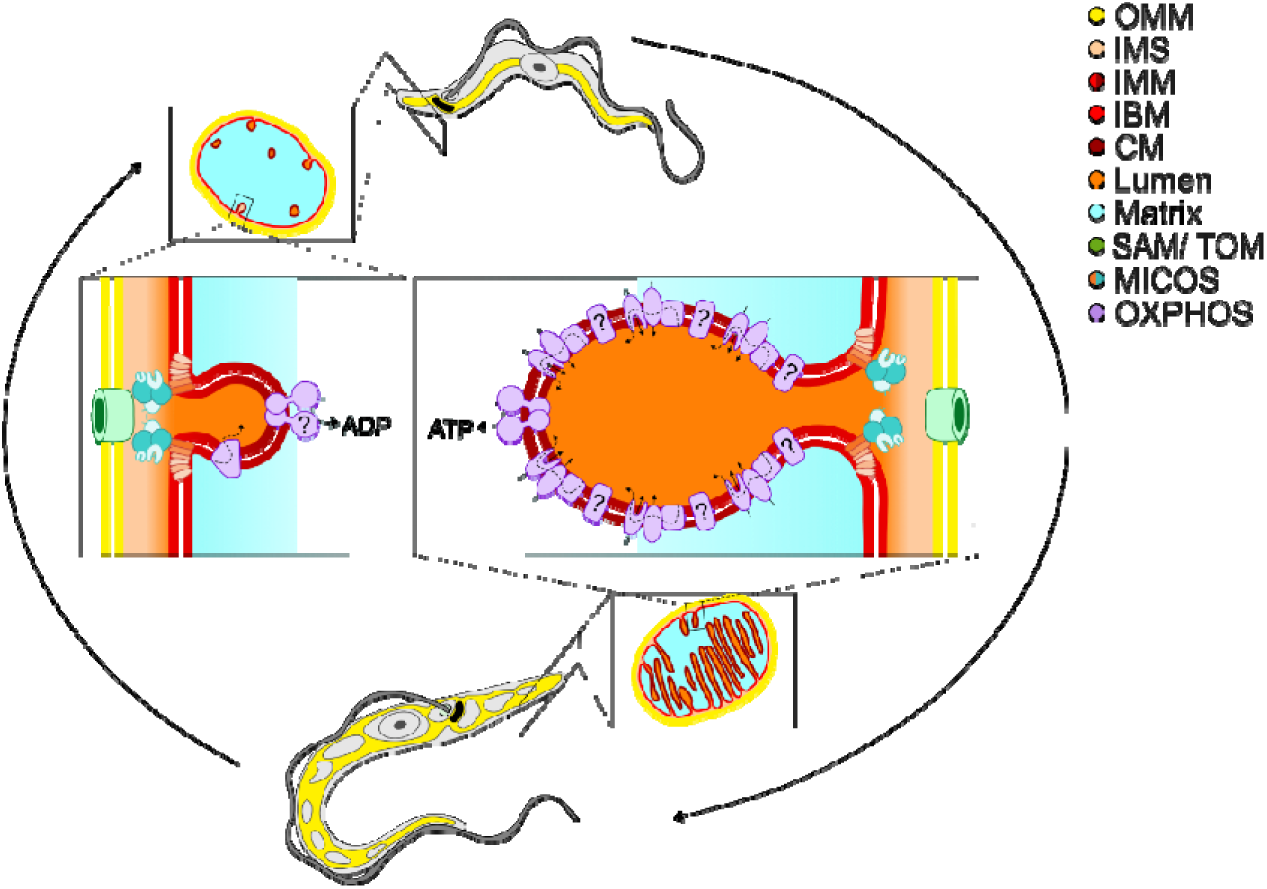
Model of mitochondrial remodeling during *T. brucei* differentiation. At the top, the bloodstream form (BSF) parasite is depicted with a tubular mitochondrion (highlighted in yellow), enlarged in the inset to show a cross-section containing sparse, stub-like precursor cristae. A selected crista is boxed and further magnified below, illustrating the absence of respiratory chain complexes and the presence of the alternative oxidase (TAO). The question mark associated with F_o_F_1_-ATP synthase indicates that localization of ATP synthase dimers to BSF precursor cristae has not yet been experimentally demonstrated. At the bottom, the procyclic form (PCF) is shown with an expanded, reticulated mitochondrial network enriched in mature discoidal cristae, as seen in the corresponding cross-section. A highlighted crista is enlarged above, showing membranes densely decorated with respiratory complexes and active oxidative phosphorylation, including ATP production by complex V. The question mark associated with complex I reflects its uncertain physiological contribution to oxidative phosphorylation in PCF *T. brucei*. This model suggests that interactions between MICOS and partner complexes, such as Sam50 and F_o_F_1_-ATP synthase, may contribute to membrane remodelling and crista maturation during differentiation. The color scheme (on the right) for mitochondrial subcompartments and protein complexes follows Fig. 1A.

In addition to these MICOS-associated factors, our datasets also identified SLP2 and pTim21 in both Mic10-2 and Mic34 IPs, suggesting a possible functional connection with MICOS. SLP2 has previously been linked to MICOS-related functions in other systems, including cristae shaping and respiratory supercomplex assembly (Mitsopoulos et al., 2015; Naha et al., 2024), while Tim21-family proteins participate in mitochondrial protein import (Chacinska et al., 2005). However, we did not detect Mic10-1 enrichment upon IP of SLP2 or pTim21. These negative results should be interpreted with caution, as pTim21 could not be successfully tagged in BSF, it therefore remains possible that their interaction with MICOS is weak, transient, indirect, or restricted to a small subpopulation of complexes, below the detection limits of our approach. Further work will be required to clarify whether SLP2 or pTim21 are functionally linked to MICOS in *T. brucei*.

The apparently more crowded molecular environment surrounding MICOS in precursor cristae may also have facilitated the detection of MIIP1 and MIIP2, two integral membrane microproteins that interact with MICOS, in both examined life cycle stages. Although we were not able to define their function, we demonstrated that their downregulation did not affect parasite growth or MICOS stability in PCF. Furthermore, both proteins share similar topology as well as predicted structure. However, only MIIP1 is conserved across kinetoplastids, whereas MIIP2 appears to have emerged in *T. brucei*. Notably, the isoelectric point (pI) of *T. brucei* MIIP1 is considerably more acidic than that of its kinetoplastid orthologs (Table S3), whereas MIIP2 exhibits an isoelectric point similar to these orthologs. Thus, MIIP2 may have arisen in *T. brucei* to compensate for the unusually acidic pI of MIIP1 in a still undefined IMM-associated function. Given that both proteins are predicted to extend about 7 nm into the IMS and that OMM-IMM contact sites are separated by ∼14 nm (Frey & Mannella, 2000), MIIP1 and/or MIIP2 could potentially form oligomeric assemblies that reinforce membrane contact sites with OMM components. Interestingly, an integral OMM protein containing an IMS-exposed coiled coil domain has recently been proposed to interact with MICOS and fortify OMM-IMM contact sites in vertebrates (Bock-Bierbaum et al., 2025), reminiscent of the hypothesized function of the MIIPs in this study. However, important differences limit this analogy: the vertebrate protein is considerably larger, includes an additional C-terminal helix bundle, and is anchored in the OMM. Future work on these and other factors will reveal how contact sites are established and maintained across different lineages.

## Materials and Methods

### Trypanosome culture and growth curves

Monomorphic BSF *T. brucei* SmOxB4 (Poon et al., 2012) were cultured in HMI-11 (Hirumi & Hirumi, 1989) supplemented with heat-inactivated (h.i.) 10% (v/v) fetal bovine serum (FBS) and 0.1 μg/ml puromycin at 37°C in the presence of 5% CO_2_. Genetically modified parasites were cultivated in HMI-11 medium in the presence of appropriate antibiotics to maintain their genetic background (2.5 μg/ml G418, 5 μg/ml hygromycin).

Pleomorphic BSF trypanosomes AnTat1.1 were cultured in HMI-11 supplemented with h.i. 10% (v/v) FBS at 37°C and 5% CO_2_. Induction of differentiation into PCF was performed as previously described (Overath et al., 1986), with slight modifications. In brief, cultures were set up in HMI-11 at ∼5 × 10L cells/ml and incubated for 48 h at 37°C and 5% CO_2_. Cells were harvested by centrifugation at 1,000 × g for 10 min and resuspended to density of 4 × 10L cells/ml in DTM medium supplemented with h.i. 15% (v/v) FBS, 7.5 μg/ml hemin, sodium citrate along with cis-aconitate were added to final concentration of 3 mM to induce differentiation. Cultures were then maintained at 27°C for 48 h, harvested by centrifugation, and resuspended in SDM80 supplemented with h.i. 10% (v/v) FBS to reach cell density of 1 × 10□/ml. Thereafter, cells were maintained in exponential growth by dilution with SDM80 supplemented with h.i. 10% (v/v) FBS.

Procyclic *T. brucei* SmOxP9 were cultured in SDM79 or SDM80 medium (Coustou et al., 2008), the latter lacking glucose, containing 1 μg/ml puromycin at 27°C. Both media were supplemented with h.i. 10% (v/v) FBS and 7.5 μg/ml hemin. Genetically modified parasites were cultivated in the presence of appropriate antibiotics to maintain their genetic background (2.5 μg/ml phleomycin, 50 μg/ml hygromcycin and 15 μg/ml G418). RNAi induction was triggered by the addition of 1 μg/ml doxycycline every 48h. Throughout all analyses, cells were maintained in exponential growth phase.

For growth curve analysis, PCF *T. brucei* cultures were grown in triplicate in SDM80 medium supplemented with h.i. 10% (v/v) FBS and appropriate antibiotics, in the presence or absence of doxycycline. Cell density was determined using the Beckman Coulter Z2 Cell and Particle Counter, cultures were diluted to 1 × 10L cells/ml every 48 h to maintain exponential growth.

### Generation of transgenic cell lines

To endogenously tag MICOS subunits with HA epitope, a modified version of pPOTv4 (Kaurov et al., 2018) was used as a template (primers listed in Table S5). PCR products were purified by phenol-chloroform extraction, ethanol-precipitated, and transfected by electroporation into BSF SmOxB4 cells using homemade transfection buffer (Schumann Burkard et al., 2011) and an Amaxa Nucleofector (Lonza), program X-001. Selection was initiated at least 8 h post-transfection using 2.5 μg/ml G418. Clones were typically apparent after 7 days, and 3 clones were analyzed by western blotting.

Potential MICOS interactors were endogenously tagged with V5 epitope using a modified version of pPOTv4 (Kaurov et al., 2018; primers listed in Table S5) in a Mic34-HA expressing background. Tagging was performed as described above, except that 5 μg/ml hygromycin was used for selection.

Tagging in PCF SmOxP9 cells was performed as described for BSF, except that a single PCR reaction was directly transfected using Amaxa Nucleofector (Lonza), program X-014. Selection antibiotics were added at least 8 h post-transfection (15 μg/ml G418 for HA-tagging and 50 μg/ml hygromycin for V5-tagging constructs).

RNAi constructs were generated by cloning a region of the coding sequence designed by RNAit (Redmond et al., 2003; primers in Table S5) into the expression vector p2T7-177 (Wickstead et al., 2002), using BamHI and XhoI restriction sites. The double RNAi construct for simultaneous downregulation of two target genes was generated using a three-way ligation with an XbaI site positioned centrally. Approximately 12 µg of each plasmid was linearized with NotI, ethanol-precipitated, and transfected into a cell line expressing MIIP1-V5 and MIIP2-HA. Selection was performed using 2.5 µg/ml phleomycin, and clones were verified by western blotting.

### Selective permeabilization with digitonin

Cells were separated into cytoplasmic and organellar fractions as previously described (Kaurov et al., 2018). Briefly, 1 × 10^8^ cells were treated with 0.015% (w/v) digitonin for 5 min on ice. Following centrifugation at 5,000 × g for 5 min at 4°C, the supernatant representing the cytosolic fraction and the pellet containing the organellar fraction were processed for further analysis by either SDS-PAGE, IP, proteinase protection assay or BN-PAGE.

### SDS-PAGE and Western Blotting

Protein samples were separated by home-made 12% SDS-PAGE containing 0.5% (v/v) 2,2,2-trichloroethanol allowing for total protein visualization *via* ultraviolet-light exposure. The proteins were then transferred onto PVDF membrane (Amersham), and probed with the appropriate primary antibodies diluted with 5% milk (w/v) in PBS supplemented with 0.1% (v/v) Tween20: Mic10-1 (1:1,000, Kaurov et al., 2018), Mic20 (1:1,000, Kaurov et al., 2022), Mic32 (1:1,000, Kaurov et al., 2022), Mic34 (1:1,000, Sheikh et al., 2025), paraflagellar rod L13D6 (1:200, Kohl et al., 1999), HA (1:1,000, Sigma), enolase (1:1,000, provided by P.A.M. Michels), Hsp70 (1:1,000, provided by A. Zíková), V5 (1:1,000, Invitrogen) and Erv1 (1:1,000, Basu et al., 2013).

Secondary HRP-conjugated anti-rabbit or anti-mouse antibodies (1:2,000, Sigma) were used. Proteins were visualized using the Pierce ECL system on a ChemiDoc instrument (Bio-Rad). Precision Plus Protein Dual Color Standards (Bio-Rad) was used as molecular weight marker.

Western Blot quantification was performed using Image Lab (Bio-Rad).

### Immunoprecipitation

Crude mitochondrial fractions were prepared from 5 × 10^8^ as described above, these were solubilized in mitochondrial lysis buffer (20 mM Tris-HCl (pH 7.4), 50 mM NaCl, 10% (v/v) glycerol, 0.1 mM EDTA, 1 mM PMSF, 1,5% (w/v) digitonin) on ice for 1 h. Following centrifugation at 16,000 × g for 30 min at 4°C, the supernatant was collected as the cleared lysate. An aliquot was retained as the input (IN) fraction, and the remaining lysate was incubated with HA-coupled paramagnetic beads (Thermo Fisher) for 2 h at 4°C with rotation. The unbound material was then collected as the flow-through (FT) fraction, beads were washed five times with lysis buffer. An aliquot of the beads was retained as the eluate (EL) for analysis by SDS-PAGE and the remaining beads were stored at −80°C until mass spectrometry analysis. All IPs were performed in triplicate.

### Mass-Spectrometry

Label-free quantification mass spectrometry analysis (LFQ-MS) of immunoprecipitates was performed by on-bead digestion. Briefly, washed beads with immunocaptured complexes were solubilized with 1% (w/v) sodium deoxycholate in 100 mM TEAB (triethylammonium bicarbonate), reduced with 5 mM TCEP [tris(2-carboxyethyl)phosphine], alkylated with 10 mM MMTS (S-methyl methanethiosulfonate), digested with Lys-C and trypsin, and extracted with ethylacetate saturated with water (Masuda et al., 2008). Tryptic peptides were desalted on Empore C18 StageTips and approximately 1 μg of peptide digest was separated on a 50 cm C18 column (EASY-Spray ES903, Thermo Fisher) using a 90 min elution gradient (Dionex Ultimate 3000, flowrate 300 nl/min) and analyzed in a DDA mode on Orbitrap Exploris 480 mass spectrometer equipped with a FAIMS unit operated at −40 and −60 V compensation voltages.

Raw files were converted to mzXML files, each containing separate compensation voltage (CV, −40, −60 V) scans using FAIMS MzXML Generator (release 1.1.8003, https://github.com/PNNL-Comp-Mass-Spec/FAIMS-MzXML-Generator) and analyzed in MaxQuant (v. 1.6.17.0, Tyanova et al., 2016). Label-free quantification (LFQ) algorithm MaxLFQ was activated and FDR was set as 0.01 at all levels. *T. brucei brucei* 927 annotated protein database (Version 58, TriTrypDB; Aslett et al., 2010) supplemented with frequently observed contaminants was used. MMTS alkylated cysteine was selected as a fixed modification (Methylthio (C), composition: H(2) C S, +45.988). Variable modifications were oxidation (M) and acetyl (protein N-term). Downstream processing of the proteinGroups.txt file was performed in Perseus (v. 1.6.10; Tyanova & Cox, 2018). Proteins were considered significantly enriched following a positive two sample student’s t-test with an FDR of 0.05 and co-enrichment across biological replicates.

### Proteinase K protection assay

The Proteinase K protection assay was performed as previously described (Kaurov et al., 2018), with slight modifications. Following selective permeabilization with digitonin, the organellar fraction was further treated with 0.05% (w/v) digitonin for 15 min on ice and centrifuged as described above to obtain mitoplasts. These were then resuspended in SoTE buffer and split into three tubes; two were treated with 10 μg proteinase K each, and one of these was additionally supplemented with 1% (v/v) Triton X-100. The remaining tube was left untreated as a mock control. All samples were incubated on ice for 15 min. Finally, the reaction was terminated by addition of 5 mM PMSF.

### BN-PAGE

Mitochondria-enriched fractions were resuspended in solubilization buffer (50 mM NaCl, 50 mM Bis-Tris, 2 mM 6-aminohexanoic acid, 1 mM EDTA, cOmplete™ Mini Protease Inhibitor Cocktail (Roche), pH 7 at 4°C), and solubilized by addition of 2% (w/v) n-Dodecyl-β-D-maltoside (Sigma) for 1 h on ice. Following centrifugation (16,000 × g, 30 min, 4°C), protein concentration was determined using Pierce™ BCA Protein Assay Kit (Thermo Fisher). Approximately 55 µg of protein was loaded onto a precast Native PAGE 3-12% Bis-Tris Gel (Invitrogen). Electrophoresis was performed at 80 V few minutes at room temperature, followed by 100 V for 3,5 h at 4°C, and final 30 min at 130 V. Proteins were blotted onto PVDF membrane (Amersham) by wet transfer (100 V, 2 h, 4°C). Membranes were blocked with 5% (w/v) milk in PBS and incubated overnight at 4°C with primary antibodies against V5 (1:1,000, Sigma, produced in rabbit), HA (1:500, Invitrogen, produced in mouse), Mic10-1 (1:500 kindly provided by A. Schneider) or ATP synthase subunit β (1:2000, provided by A. Zíková). Secondary antibody incubation and visualization was performed as described for SDS-PAGE.

### Immunofluorescence Assay

Cells were pelleted, washed once with PBS and spread onto microscope slides. After air-drying at room temperature (RT), samples were fixed in ice-cold methanol for 10 min. The slides were rehydrated in PBS and blocked with 3% (w/v) BSA in PBS for 30 minutes at RT. Primary antibodies were applied for 1 h at RT as follows: high affinity HA (1:100, Roche), V5 (1:100, Invitrogen, produced in mouse) and Hsp70 (1:500, provided by A. Zíková). Slides were washed twice with PBS and incubated for 1 h at RT in the dark with appropriate secondary antibodies. Goat anti-rat Alexa Fluor 488 and goat anti-mouse Alexa Fluor 555 (1:1000, Invitrogen) were used for detection of HA and Hsp70 signals, respectively, while goat anti-mouse Alexa Fluor 488 (1:2000, Invitrogen) was used to visualize the V5 signal. Samples were mounted in ProLong antifade reagent containing DAPI, sealed, and stored at 4°C until imaging with Olympus BX63 widefield fluorescence microscope. For MitoTracker Red staining (Invitrogen), cells were incubated with 10 μM dye for 30 minutes under standard culture conditions prior to processing as described above.

### Immunolectron Microscopy

For immunogold labeling, cells were fixed with 4% (w/v) formaldehyde and 0.1% (v/v) glutaraldehyde in 0.1 M HEPES for 1 h at RT. After washing, pellets were embedded in 10% (w/v) gelatin, immersed in 2.3 M sucrose for 24 h at 4°C, and frozen by plunging into liquid nitrogen. Cryosections were cut using EM UC6 ultramicrotome (Leica). Cryosections were picked up with a drop of 1.15 M sucrose/1% (w/v) methylcellulose. The sections were incubated in blocking solution (2% (w/v) BSA, 0.05% (v/v) Tween20 in HEPES) for 1 h at RT. The sections were immunolabeled with rat anti-HA antibody (Roche) diluted 1:20. After washing with blocking solution, the samples were incubated for 1 h with protein A conjugated to 10 nm gold nanoparticles (UMC, Utrecht) diluted 1:50. Samples were washed in HEPES and dH_2_O, contrasted, and embedded in 1.8% (w/v) methylcellulose/0.3% (w/v) uranyl acetate.

Samples were observed with a JEOL 1400 transmission electron microscope (TEM) operating at an accelerating voltage 120 kV equipped with Xrosa CMOS camera (EMSIS). Electron tomograms were collected using the JEOL 2100F TEM at 200 kV, controlled with SerialEM automated acquisition software, equipped with a high-tilt stage and direct electron detector K2 Summit (Gatan). Tomograms were reconstructed using the IMOD software package. Crista membranes were enhanced using the IMODs nonlinear anisotropy diffusion filter (K-value, 30; iterations, 20) to compensate for their low contrast staining due to Tokuyasu processing for immunogold labelling (Tokuyasu, 1978).

### In silico analysis

Kinetoplastid orthologs were searched by BLAST in TriTrypDB (Aslett et al., 2010) using *T. brucei* pTim21, MIIP1, and MIIP2 as queries against the kinetoplastid genomes listed in Tables S3-S4. pTim21 structural homology searches were performed using HHPred (Zimmermann et al., 2018) and FoldSeek (van Kempen et al., 2024). In the former case, a trimmed multisequence MAFFT (Katoh et al., 2019) alignment (MSA) of pTim21 was used as the query, whereas the *T. brucei* pTim21 AlphaFold structure (accession number Q384E9) was used to search FoldSeek. All HHPred hits above 95% probability and FoldSeek hits to entries in the SwissProt database with 100% probability are shown in Fig S3. HHPred search queried the following databases: PDB_mmCIF70_25_May, Pfam-A_v37.4, SMART_v6.0, TIGRFAMs_v15.0. Searches were performed on 17 November 2025.

MIIP1 MSA shown in Fig. S6C was used as a query for JackHMMer (Potter et al., 2018), in which homology to the t-SNARE CC domain was detected after two iterations (Figs. 5A, S6D). CC and TMD prediction were performed using Coils2 (Lupas et al., 1991) and TMHMM (Krogh et al., 2001) implemented in Geneious software (Dotmatics). Mitochondrial targeting was predicted using DeepMito (Savojardo et al., 2020) or PSORT II (https://psort.hgc.jp/form2.html). AlphaFold3 (Abramson et al., 2024) was used to predict MIIP monomeric and dimeric structures, which were visualized using ChimeraX (Meng et al., 2023).

## Acknowledgements

We thank Shaghayegh ‘Nina’ Sheikh for her valuable input and advice, Julius Lukeš for his support and Sofija Hitac for help with some experiments. We acknowledge the Proteomics Service Laboratory at the Institute of Physiology (supported by RVO, ID 67985823) and the Institute of Molecular Genetics (supported by RVO, ID 68378050) of the Czech Academy of Sciences for mass spectrometry-based proteomic analyses, with special thanks to Marek Vrbacký. This study was supported by Czech Science Foundation (Grant 23-07674S to HH), USB Grant Agency (Grant 121/2024/P to MB, 04-070/2025/P) and the BC CAS core facility LEM supported by MEYS CR (LM2023050 Czech-BioImaging and OP VVV CZ.02.1.01/0.0/0.0/18_046/0016045).

## Author Contributions

MB, CB and HH, Conceptualization; MB, CB, TB and HH, Formal Analysis; MB, TW, CB, TB, MT, and HH, Investigation; CB and HH, Supervision; MB, CB, TB, and HH, Visualization; MB, CB, and HH, Writing – Original Draft Preparation; MB, CB, TB and HH, Writing – Review & Editing; HH, Funding Acquisition.

## Data availability

The mass spectrometry proteomics data have been deposited to the ProteomeXchange Consortium via the PRIDE partner repository (Perez-Riverol et al., 2025) with the dataset identifier PXD074741.

## Ethics statement

No ethical was approval required.

## Copyright statement

No copyright permission needed to be taken.

## Supplemental material

**Fig. S1. Generation of bloodstream form *T. brucei* cell lines expressing C-terminally HA-tagged MICOS subunits. (A)** Immunoblot analysis confirming successful HA-tagging of the indicated MICOS subunits (above the blot). Molecular weight is shown on the left and antibody on the right. Immunodetection of Hsp70 served as a loading control (Load). **(B)** Immunofluorescence assay of the indicated cell lines. Antibodies are indicated above the panel; Hsp70 marks mitochondria, and DAPI (blue) stains DNA. The composite image demonstrates substantial overlap of the fluorescence signals. Scale bar, 5 μm.

**Fig. S2. Z-projection of tomograms shown in Fig. 2C. (A)** First tomogram from the left in Fig. 2C. Top left image is the Z-projection, with boxed region enlarged 2-fold below. Solid arrowheads, mitochondrial membrane; hollow arrowheads, crista membrane. Right panel show the same images with mitochondrial membrane traced in red dashed line and crista membrane in orange. See Movie S3. **(B)** As in (A), with the boxed region enlarged 3-fold. See Movie S4.

**Fig. S3. Structural homology search results for putative Tim21 (pTim21). (A)** Visualization along with hitlist generated by HHPred and **(B)** top hits identified by FoldSeek. The top four (HHPred) and top seven (FoldSeek) hits are shown, along with the next highest-ranking hit displayed at the bottom of each list. Notably, Coa1 is a homolog of Tim21 (Shinde et al., 2021). Color coding in (A): Red, ≥95% confidence; Cyan, 20-50% confidence; Blue, <20% confidence (Zimmermann et al., 2018). Only red hits were considered.

**Fig S4. Neither SLP2 nor pTim21 interacts with MICOS in *T. brucei* PCF.** Immunoblot analysis of reciprocal IPs for two candidate interactors identified in Fig. 3C-D: SLP2-V5 and pTim21-V5. Input (IN), flow-through (FT), and eluate (EL) fractions were analyzed as in Fig. 3B. Cell lines and fractions indicated above the blot, molecular weight on the left, and antibodies on the right.

**Fig. S5. Generation of V5-tagged cell lines and growth analysis following depletion of MICOS-interacting integral microproteins (MIIPs). (A)** Generation of *T. brucei* procyclic form cell lines expressing C-terminally V5-tagged MIIP1 or MIIP2, used in the experiment shown in Fig. 4D. Immunoblot analysis confirming successful tagging of the indicated proteins (shown above the blot). Molecular weight marker indicated on the left, antibody on the right. Immunodetection of Hsp70 served as a loading control (Load). **(B)** Growth analysis following inducible depletion of MIIPs. Proliferation of *T. brucei* PCF cell line carrying inducible RNAi targeting both *miip* transcripts was monitored in glucose-poor SDM80 medium. The non-induced RNAi line served as a control. Cell density was measured over time and plotted on logarithmic scale (y-axis); x-axis, days post-RNAi induction. Data represent the mean (n=3); error bars indicating standard deviation are not visible at this scale.

**Fig. S6. Multi-sequence alignment (MSA) of 13 kinetoplastid MIIP1 orthologs. (A)** Consensus sequence of aligned orthologs; amino acid coverage at each position (scale 1-13); **(B)** Mean hydrophobicity (HP) and isoelectric point (pI) profiles (as in Fig. 5A); and **(C)** sequence logo illustrating residue conservation across the MSA below. **(D)** Identified domains (as in Fig. 5A) are indicated, along with predicted coiled-coil regions in each species.

**Movie S1. Localization of Mic34-HA on precursor cristae by immunoelectron tomography.** Related to the first tomogram from the left in Fig. 2C. The mitochondrial membranes are traced in red, the crista membrane in orange, and the gold nanoparticle bound to the HA antibody is shown as a green dot, using the same color scheme as in Fig. 2C.

**Movie S2. Localization of Mic34-HA on precursor cristae by immunoelectron tomography.** Related to the third tomogram from the left in Fig. 2C. The mitochondrial membranes are traced in red, the crista membrane in orange, and gold nanoparticle bound to the HA antibody is shown as a green dot, using the same color scheme as in Fig. 2C.

**Movie S3. Virtual sections of the first tomogram from left shown in Fig 2C and S2.** The top 21 virtual sections, corresponding to a thickness of approximately 14 nm, are shown. Yellow arrows point to crista membrane.

**Movie S4. Virtual sections of the second tomogram from left shown in Fig 2C and S3.** The top 21 virtual sections, corresponding to a thickness of approximately 14 nm, are shown. Yellow arrows point to crista membrane.

**Table S1: Significantly enriched ATP synthase subunits and mitochondrial carrier proteins (MCPs) identified in Mic34-HA IP.**

**Table S2: Novel MICOS-interacting candidates significantly enriched in Mic10-2-HA and Mic34-HA IPs.**

**Table S3: Empirical and predicted localization of MICOS-interacting integral proteins (MIIP) 1 and 2 plus transmembrane domain and coiled coil prediction.** Results of N-terminal and C-terminal tagging retrieved from TrypTag database (Dean et al., 2017). PSORTII ‘NiTerm’ denotes the predicted length of the mitochondrial targeting presequence in amino acid residues; “score” indicates percentage confidence of prediction. PSORTII ‘PSG’ denotes signal peptide prediction score (positive values indicate likelihood); ‘GAVEL’, predicted cleavage site of mitochondrial presequence (|). ‘Start’, 1st amino acid coordinate relative to the start methionine; ‘End’, refers to the last amino acid coordinate relative to start methionine.

**Table S4: Kinetoplastid orthologs of putative Tim21 (pTim21) analyzed in this study.**

**Table S5: Oligonucleotides used for generation of the transgenic cell lines described in this study.**

**Table S6: Detailed mass spectrometry results Mic34-HA IP. Table S7: Detailed mass spectrometry results Mic10-2-HA IP.**

