## Supplementary figures for "Trypanosomal MICOS is assembled on non-respiring mitochondrial crista precursors and associates with two integral microproteins"

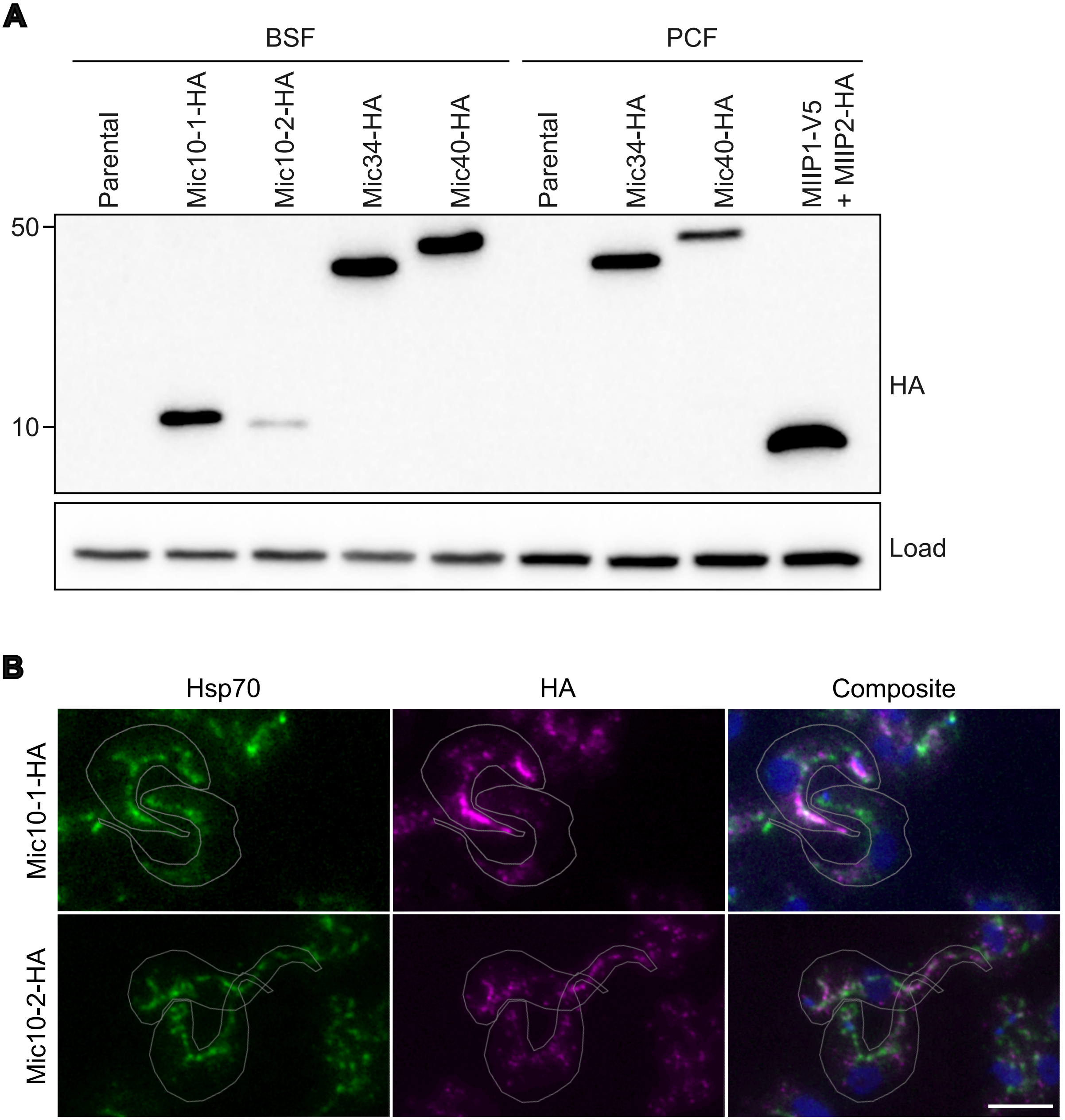


**Fig. S1. Generation of bloodstream form *T. brucei* cell lines expressing C-terminally HA-tagged MICOS subunits. (A)** Immunoblot analysis confirming successful HA-tagging of the indicated MICOS subunits (above the blot). Molecular weight is shown on the left and antibody on the right. Immunodetection of Hsp70 served as a loading control (Load). **(B)** Immunofluorescence assay of the indicated cell lines. Antibodies are indicated above the panel; Hsp70 marks mitochondria, and DAPI (blue) stains DNA. The composite image demonstrates substantial overlap of the fluorescence signals. Scale bar, 5 μm.

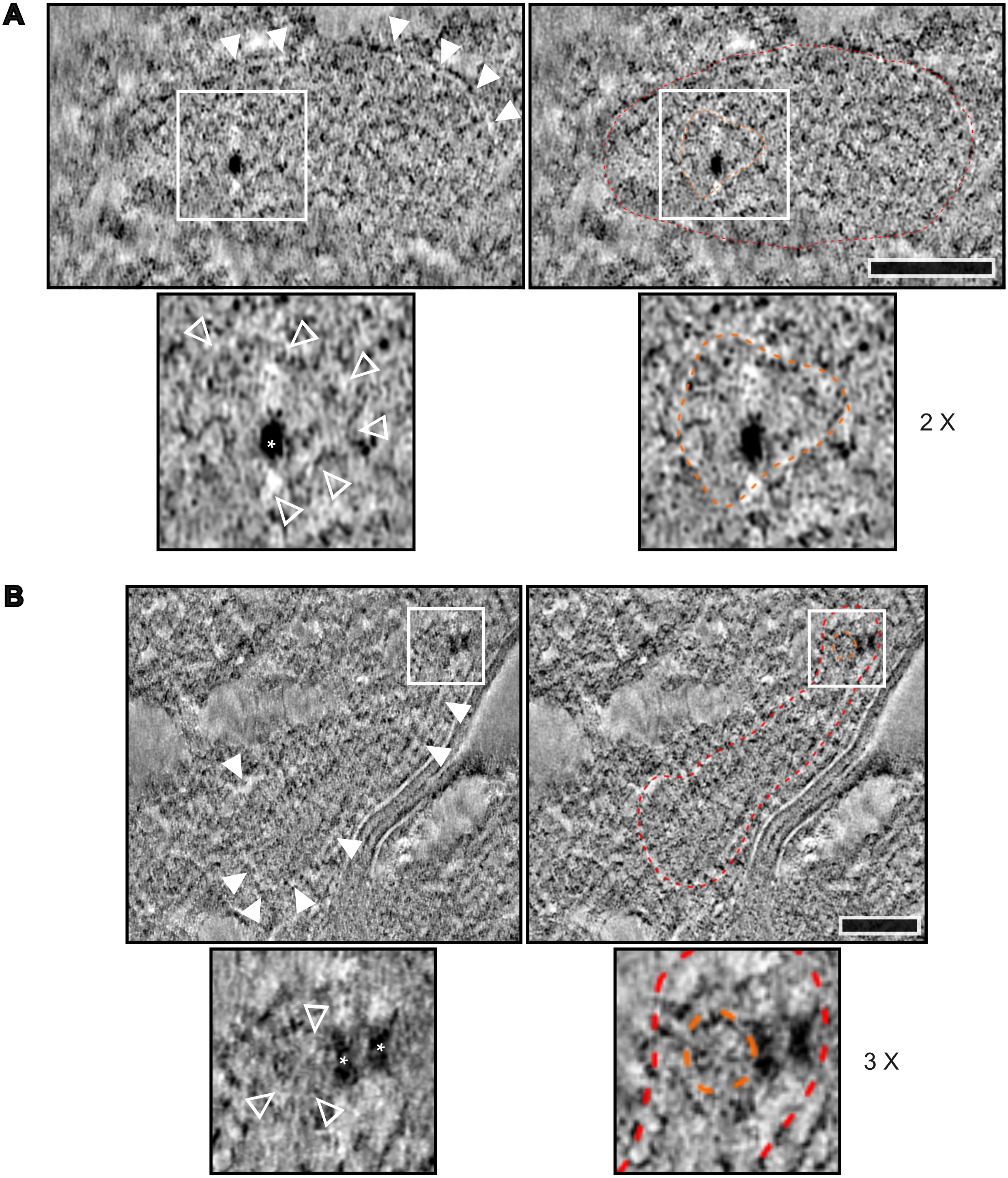


**Fig. S2 Z-projection of tomograms shown in Fig. 2C. (A)** First tomogram from the left in Fig. 2C. Top left image is the Z-projection, with boxed region enlarged 2-fold below. Solid arrowheads, mitochondrial membrane; hollow arrowheads, crista membrane. Right panel show the same images with mitochondrial membrane traced in red dashed line and crista membrane in orange. See Movie S3. **(B)** As in (A), with the boxed region enlarged 3-fold. See Movie S4.

**A**

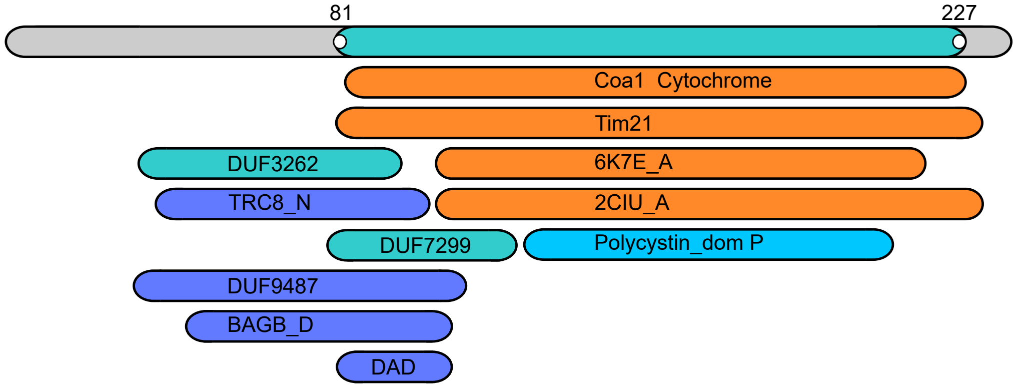


| Nr | Hit | Name | Probability | E-value | Score | SS | Aligned cols | Target length |
|---|---|---|---|---|---|---|---|---|
| 1 | PF08695.16 | Coa1; Cytochrome oxidase complex assembly protein 1 | 97.25 | 0.02 | 42.34 | 10.7 | 114 | 116 |
| 2 | PF08294.17 | TIM21 | 96.54 | 0.11 | 41.43 | 10.5 | 134 | 153 |
| 3 | 6K7E_A | Maltose/ maltodextrin-binding periplasmic protein; Mitochondrial import inner membrane translocase subunit TIM21 | 96.04 | 0.05 | 48.15 | 6.5 | 105 | 507 |
| 4 | 2CIU_A | Import inner membrane translocase subunit TIM21 | 95.53 | 0.26 | 39.30 | 8.3 | 111 | 127 |
| 5 | PF11660.13 | DUF3262; Protein of unknown function | 60.06 | 130.00 | 22.94 | 7.3 | 60 | 76 |

**B**

| Target | Description | Scientific Name | Prob. | Seq. Id. | E-value | Position in query |
|---|---|---|---|---|---|---|
| AF-Q9BVV7-F1-model_v6 | Mitochondrial import inner membrane translocase | *Homo sapiens* | 1 | 15.3 | 3.05E-05 | 1 → 264 |
| AF-Q3SZV6-F1-model_v6 | Mitochondrial import inner membrane translocase | *Bos taurus* | 1 | 16.6 | 7.99E-05 | 1 → 264 |
| AF-Q75CX4-F1-model_v6 | Mitochondrial import inner membrane translocase | *Eremothecium gossypii* | 1 | 16.3 | 5.69E-05 | 4 → 277 |
| AF-Q8CCM6-F1-model_v6 | Mitochondrial import inner membrane translocase | *Mus musculus* | 1 | 16.2 | 2.49E-04 | 1 → 264 |
| AF-Q6FMZ2-F1-model_v6 | Mitochondrial import inner membrane translocase | *Nakaseomyces glabratus* | 1 | 15.5 | 4.39E-04 | 16 → 270 |
| AF-Q6CW96-F1-model_v6 | Mitochondrial import inner membrane translocase | *Kluyveromyces lactis* | 1 | 16.4 | 2.95E-04 | 1 → 270 |
| AF-Q5U2X7-F1-model_v6 | Mitochondrial import inner membrane translocase | *Rattus norvegicus* | 1 | 15.8 | 3.07E-04 | 1 → 264 |
| AF-Q1T7B7-F1-model_v6 | Centromere protein P | *Gallus gallus* | 0.93 | 5.9 | 2.02E-02 | 74 → 255 |

**Fig. S3. Structural homology search results for putative Tim21 (pTim21). (A)** Visualization along with hitlist generated by HHPred and **(B)** top hits identified by FoldSeek. The top four (HHPred) and top seven (FoldSeek) hits are shown, along with the next highest-ranking hit displayed at the bottom of each list. Notably, Coa1 is a homolog of Tim21 (Shinde et al., 2021). Color coding in (A): Red, ≥95% confidence; Cyan, 20-50% confidence; Blue, <20% confidence (Zimmermann et al., 2018). Only red hits were considered.

**S4**

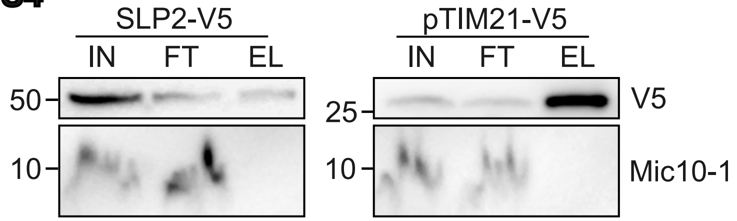


**Fig S4. Neither SLP2 nor pTim21 interacts with MICOS in *T. brucei* PCF.** Immunoblot analysis of reciprocal IPs for two candidate interactors identified in Fig. 3C-D: SLP2-V5 and pTim21-V5. Input (IN), flow-through (FT), and eluate (EL) fractions were analyzed as in Fig. 3B. Cell lines and fractions indicated above the blot, molecular weight on the left, and antibodies on the right.

**A**

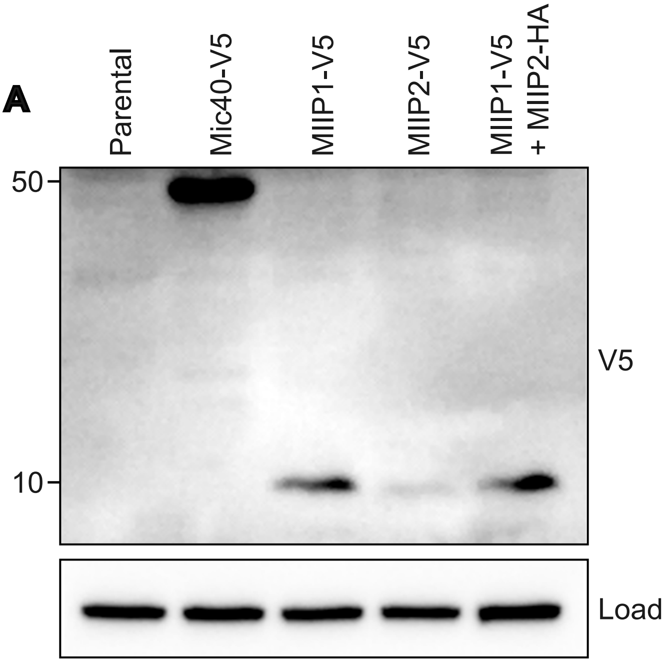


**B**

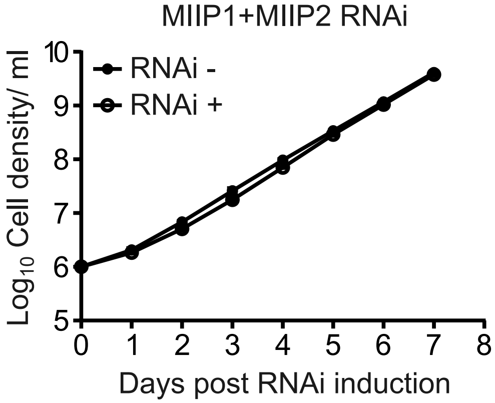


**Fig. S5. Generation of V5-tagged cell lines and growth analysis following depletion of MICOS-interacting integral microproteins (MIIPs). (A)** Generation of *T. brucei* procyclic form cell lines expressing C-terminally V5-tagged MIIP1 or MIIP2, used in the experiment shown in Fig. 4D. Immunoblot analysis confirming successful tagging of the indicated proteins (shown above the blot). Molecular weight marker indicated on the left, antibody on the right. Immunodetection of Hsp70 served as a loading control (Load). **(B)** Growth analysis following inducible depletion of MIIPs. Proliferation of *T. brucei* PCF cell line carrying inducible RNAi targeting both *miip* transcripts was monitored in glucose-poor SDM80 medium. The non-induced RNAi line served as a control. Cell density was measured over time and plotted on logarithmic scale (y-axis); x-axis, days post-RNAi induction. Data represent the mean (n=3); error bars indicating standard deviation are not visible at this scale.

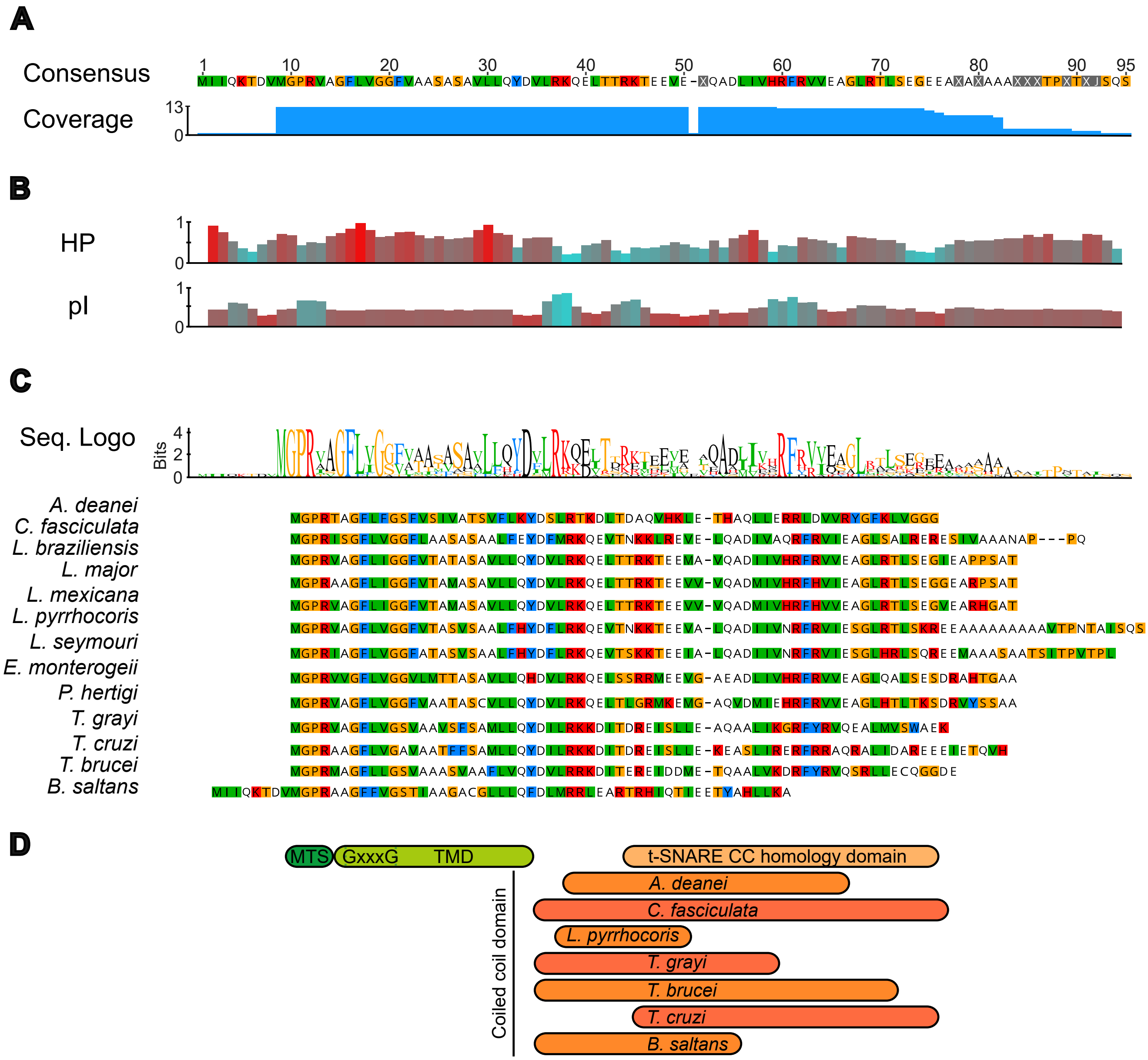


**Fig. S6. Multi-sequence alignment (MSA) of 13 kinetoplastid MIIP1 orthologs. (A)** Consensus sequence of aligned orthologs; amino acid coverage at each position (scale 1-13); **(B)** Mean hydrophobicity (HP) and isoelectric point (pI) profiles (as in Fig. 5A); and **(C)** sequence logo illustrating residue conservation across the MSA below. **(D)** Identified domains (as in Fig. 5A) are indicated, along with predicted coiled-coil regions in each species.

**Movie S1. Localization of Mic34-HA on precursor cristae by immunoelectron tomography.** Related to the first tomogram from the left in Fig. 2C. The mitochondrial membranes are traced in red, the crista membrane in orange, and the gold nanoparticle bound to the HA antibody is shown as a green dot, using the same color scheme as in Fig. 2C.

**Movie S2. Localization of Mic34-HA on precursor cristae by immunoelectron tomography.** Related to the third tomogram from the left in Fig. 2C. The mitochondrial membranes are traced in red, the crista membrane in orange, and gold nanoparticle bound to the HA antibody is shown as a green dot, using the same color scheme as in Fig. 2C.

**Movie S3. Virtual sections of the first tomogram from left shown in Fig 2C and S2.** The top 21 virtual sections, corresponding to a thickness of approximately 14 nm, are shown. Yellow arrows point to crista membrane.

**Movie S4. Virtual sections of the second tomogram from left shown in Fig 2C and S3.** The top 21 virtual sections, corresponding to a thickness of approximately 14 nm, are shown. Yellow arrows point to crista membrane.
